# *Ppd-H1* integrates drought stress signals to control spike development and flowering time in barley

**DOI:** 10.1101/2020.03.26.010173

**Authors:** Leonard Gol, Einar B. Haraldsson, Maria von Korff

**Author notes:** Correspondence: Maria von Korff. Author Contact Information: Leonard Gol, Einar B. Haraldsson.

## Abstract

Drought impairs growth and spike development and is therefore a major cause of yield losses in the temperate cereals barley and wheat. Here, we show that the photoperiod response gene *PHOTOPERIOD-H1* (*Ppd-H1*) interacts with drought stress signals to modulate spike development. We tested the effects of a continuous mild and a transient severe drought stress on developmental timing and spike development in spring barley cultivars with a natural mutation in *ppd-H1* and derived introgression lines carrying the wild-type *Ppd-H1* allele from wild barley. Mild drought reduced the spikelet number and delayed floral development in spring cultivars but not the introgression lines with a wild-type *Ppd-H1* allele. Similarly, drought-triggered reductions in plant height, tiller and spike number were more pronounced in the parental lines compared to the introgression lines. Transient severe stress halted growth and floral development, upon rewatering introgression lines, but not the spring cultivars, accelerated development so that control and stressed plants flowered almost simultaneously. These genetic differences in development were correlated with a differential downregulation of the flowering promotors *FLOWERING LOCUS T1* and the BARLEY MADS*-*box genes *BM3* and *BM8.* Our findings, therefore, demonstrate that *Ppd-H1* affects developmental plasticity in response to drought in barley.

**Highlight:** We show that *Ppd-H1* integrates photoperiod and drought stress signals via *FLOWERING LOCUS T* 1 (*FT1)* and the downstream MADS-box genes *BM3* and *BM8* to modulate reproductive development, and shoot and spike morphology in barley.

## 1 Introduction

Global warming increases the frequency and intensity of severe water scarcity events, which negatively affect the yield of rain-fed crops such as barley (*Hordeum vulgare* L.) and wheat (*Triticum aestivum* L.) (Xie *et al.*, 2018; Kahiluoto *et al.*, 2019). Drought during reproductive development impairs spike development and floret fertility and is therefore a major cause of yield losses in these temperate cereals (Gol *et al.*, 2017). At present, strategies to breed cereal varieties with improved yield under drought are limited due to a lack of knowledge on the genetic factors that control inflorescence and flower development under drought conditions. Understanding the plasticity and genetic control of stress-induced changes in reproductive development will be crucial to ensure future yield stability of temperate cereals.

The model plant *Arabidopsis thaliana* accelerates reproductive development under drought, a response that has been termed drought escape. In Arabidopsis drought escape is triggered under inductive long day (LD) conditions and is controlled by components of the circadian clock and the photoperiod response pathway (Riboni *et al.*, 2013, 2016). Under drought conditions the phytohormone abscisic acid (ABA) modulates the activity and signalling of the clock gene *GIGANTEA* (*GI*) and consequently its ability to activate *FLOWERING LOCUS T* (*FT*) under long photoperiods (Riboni *et al.*, 2013, 2016). The FT protein acts as a florigenic signal, moving long distance from the leaf to the shoot apical meristem (SAM) to induce the floral transition (Abe *et al.*, 2005; Wigge *et al.*, 2005; Corbesier *et al.*, 2007; Tamaki *et al.*, 2007; Jaeger and Wigge, 2007; Mathieu *et al.*, 2007; Jaeger *et al.*, 2013). Under non-inductive short days (SD), ABA delays flowering by repressing the flowering-promoting MADS-box gene *SUPPRESSOR OF OVEREXPRESSION OF CONSTANS1* (*SOC1*), a transcription factor integrating floral cues in the shoot meristem (Riboni *et al.*, 2016). In addition, it was shown that ABA-responsive element (ABRE)-binding factors (ABFs) interact with NUCLEAR FACTOR Y subunit C (NF-YC) 3/4/9 to promote flowering by inducing *SOC1* transcription under drought conditions (Hwang *et al.*, 2019). On the other hand, ABSCISIC ACID-INSENSITIVE 3/4/5 bZIP transcription factors involved in ABA signalling repress flowering by upregulating the floral repressor and vernalisation gene *FLOWERING LOCUS C* (*FLC*) (Wang *et al.*, 2013; Shu *et al.*, 2016). Consequently, drought cues depend on the photoperiod and interact with photoperiod response and vernalisation genes to modulate flowering time in Arabidopsis. In contrast to Arabidopsis, rice (*Oryza sativa* L.) shows a delay in flowering in response to drought under inductive photoperiods and this delay is accompanied by a downregulation of the florigenic signals *HEADING DATE 3a* (*Hd3a*) and *RICE FLOWERING LOCUS T 1* (*RFT1*) (Zhang *et al.*, 2016; Galbiati *et al.*, 2016). Consequently, the developmental response to drought varies within and between species and is linked to the differential regulation of *FT*-like genes (Kazan and Lyons, 2016). However, the effects of drought on reproductive development and genetic components that modulate this response are not known in most crop species including the important temperate crop barley.

Barley germplasm is characterised by high genetic diversity and variation in response to abiotic stresses. While elite cultivars tend to be more stress susceptible, wild and landrace barley genotypes are well-adapted to drought-prone environments and therefore represent a valuable resource for improving stress tolerance in elite barley (Baum *et al.*, 2007; von Korff *et al.*, 2008; Rollins *et al.*, 2013*b*; Templer *et al.*, 2017). It was demonstrated that yield stability in the field was associated with the major photoperiod response gene *PHOTOPERIOD H1* (*Ppd-H1*) and the vernalisation gene *VERNALIZATION 1* (*VRN1*) (von Korff *et al.*, 2008; Rollins *et al.*, 2013*a*; Al-Ajlouni *et al.*, 2016; Wiegmann *et al.*, 2019). These findings suggested that the timing of reproductive development is crucial to maximise yield formation under harsh environmental conditions. However, it is not known if and how these floral regulators interact with stress cues to modulate development. *Ppd-H1*, a barley homolog of the PSEUDO RESPONSE REGULATOR (PRR) genes from the Arabidopsis circadian clock, induces the expression of *FLOWERING LOCUS T1* (*FT1)*, a homolog of Arabidopsis *FT* and rice *Hd3a* under LDs (Turner *et al.*, 2005; Corbesier *et al.*, 2007; Tamaki *et al.*, 2007; Campoli *et al.*, 2012*b,a*). In barley, the upregulation of *FT1* in the leaf is correlated with induction of the MADS-box genes *VRN1 (BM5a), BARLEY MADS-box 3* (*BM3*) and *BM8*, barley homologs of Arabidopsis *APETALA1/FRUITFUL* (*AP1/FUL*), and the acceleration of inflorescence development (Schmitz *et al.*, 2000; Trevaskis *et al.*, 2007; Digel *et al.*, 2015). Homologs of *Ppd-H1*/*PRR37* function in the circadian clock in Arabidopsis and rice (Makino *et al.*, 2001; Murakami *et al.*, 2003; Turner *et al.*, 2005). The circadian clock is an internal timekeeper that allows plants to anticipate predictable changes in the environment and controls a number of output traits including development and stress responses (Sanchez *et al.*, 2011; Müller *et al.*, 2014; Johansson and Staiger, 2015). In Arabidopsis, the central oscillator is composed of negative transcriptional feedback loops: the rise of *CIRCADIAN CLOCK ASSOCIATED1* (*CCA1*) and *LATE ELONGATED HYPOCOTYL* (*LHY*) late at night inhibit the evening complex genes *EARLY FLOWERING 3* (*ELF3*), *EARLY FLOWERING 4* (*ELF4*) and *LUX* which in turn repress the *PRRs* at night. Barley homologs of these clock genes have been identified and their interactions are largely conserved in barley (Campoli *et al.*, 2012*b*; Müller *et al.*, 2019). Accordingly, elements of the evening complex genes repress *Ppd-H1* at night and thereby control the photoperiod dependent upregulation of *FT1* (Mizuno *et al.*, 2012; Zakhrabekova *et al.*, 2012; Faure *et al.*, 2012; Campoli *et al.*, 2013; Alvarez *et al.*, 2016). In spring barley grown in northern latitudes, a recessive mutation in the CONSTANS, CONSTANS-like, and TOC1 (CCT) domain of *ppd-H1* has been selected (Jones *et al.*, 2008). This *ppd-H1* allele delays flowering under LDs and thereby improves yield in temperate environments with long growing seasons (Cockram *et al.*, 2007; Alqudah *et al.*, 2014; Digel *et al.*, 2015). By contrast, early flowering in response to LDs promoted by the wild-type *Ppd-H1* allele was associated with improved yield under Mediterranean environments with terminal stress (Wiegmann *et al.*, 2019). However, it is not known if the two *Ppd-H1* variants also interact with stress cues to modulate reproductive development.

Here, we provide a detailed analysis of barley development under drought. We show that variation at *Ppd-H1* interacts with drought to control flowering time, grain yield, as well as the expression of *FT1* and the downstream MADS-box genes *BM3* and *BM8*.

## 2 Materials and Methods

### 2.1 Plant materials, growth conditions and phenotyping

Drought responses were scored in the spring barley (*Hordeum vulgare* L.) cultivars Scarlett, Golden Promise and Bowman and their derived introgression lines S42-IL107 (Scarlett), GP-fast (Golden Promise) and BW281 (Bowman). Scarlett, Golden Promise and Bowman carry a natural mutation in the CCT domain of *Ppd-H1*, that causes a delay in flowering under LD conditions (Turner *et al.*, 2005). The derived introgression lines S42-IL107 and BW281 carry a dominant *Ppd-H1* allele introgressed from wild and winter barley, respectively (Druka *et al.*, 2011; Schmalenbach *et al.*, 2011). GP-fast was created via crossing of Golden Promise to the winter barley cultivar Igri, followed by two rounds of backcrossing to Golden Promise to reduce the size of the introgression.

The three spring barley cultivars and derived introgression lines were genotyped with the Barley 50k iSelect SNP Array at TraitGenetics GmbH (Gatersleben, Germany) (Bayer *et al.*, 2017). Chromosomal positions for each marker were obtained from the POPSEQ_2017 genetic map (Cantalapiedra *et al.*, 2015; Mascher *et al.*, 2017). Sizes of the introgressions were calculated based on half the distance between the markers flanking donor introgressions and the first polymorphic markers within the introgressions (Supplemental Figure **1**, Supplemental Data 1).

We conducted two different drought experiments. First, a continuous drought treatment was applied by a controlled dry down of the soil to a soil water content (SWC) of 15% of field capacity (FC) and this FC was maintained until plant maturity. In a second experiment, a transient drought treatment was applied by withholding water for eight consecutive days during floral development followed by rewatering to control levels. Both experiments were performed in a controlled environment chamber under 60% relative humidity. Grains were sown in 7 cm × 7 cm × 8 cm black plastic pots; 40 pots (5 × 8 rows) per tray. Each pot was filled with exactly 150 g of soil mixture. A mixture of 93% (v/v) Einheitserde ED73 (Einheitserde Werkverband e.V., Sinntal_Altengronau, Germany), 6.6% (v/v) sand and 0.4% (v/v) Osmocote exact standard 3–4M (Scotts Company LLC), was freshly prepared before sowing. This porous soil mixture with high organic matter content was selected to further aid the even distribution of moisture in the soil. Grains were stratified in well-watered soil at 4°C in the dark for at least four days. Plants were then germinated under SD conditions (8 h, 22°C day; 16 h, 18°C night; photosynthetically active radiation ≈250 µM/m^2^*s) For the continuous drought treatment water was withheld after germination until the SWC reached 15% FC while the control plants were watered to maintain 70% of FC. The desired SWC of 15% FC was reached after ten days when all plants were transferred from SD to LD and kept under LD for the rest of the experiment (16 h, 22°C day; 8 h, 18°C night; photosynthetically active radiation ≈250 µM/m^2^*s). For the application of severe drought, plants of Scarlett and S42-IL107 were germinated under SD conditions and shifted to LDs after ten days. All plants were kept at 70% FC until they had reached the awn primordium stage (W3). Then watering was stopped for eight consecutive days. SWC in the pots reached a relative water content of 8% FC on the eighth day. Control plants were kept at 70% FC during this time. Subsequently, all drought-treated pots were rewatered to control levels of 70% FC. FC was calculated from the difference in weight of fully hydrated and oven dried soil. SWC was measured gravimetrically (Coleman, 1947). Pots were soaked with water and subsequently left to drain by gravity until their weight remained stable; this was set as 100% FC. Dry weight was measured after pots were dried in a drying cabinet at approximately 60°C until their weight remained stable. Measurements of FC were corrected for the biomass accumulation of growing plants as the experiments progressed. The weight of pots was checked daily and all plants were watered daily to maintain the same SWC throughout development. At least three replicate plants of all six genotypes were sown and germinated for each sampling time point.

The development of the main shoot apex (MSA) was scored in accordance to the stages described by Waddington *et al.* (1983) that is based on the progression of inflorescence initiation and then the most advanced floret primordium and pistil of the inflorescence. At Waddington stage 2 (W2) the first spikelets initiate and the MSA transitions to a reproductive inflorescence. The first floral organ primordia differentiate and stem elongation initiates at the stamen primordium stage (W3.5). New spikelet primordia are continuously initiated until about W5, which then mature into florets until anthesis and pollination at W10. MSA dissection was performed with microsurgical stab knives ([SSC#72-1551], Sharpoint, Surgical Specialties Corporation). Images of developing apices were obtained using a Nikon stereo microscope (Nikon SMZ18), Nikon DS-U3 controller unit and a Nikon DS-Fi2 digital camera. Nikon NIS-Elements software was used for image acquisition. Heading date was scored at Zadoks stage Z49 when first awns became visible (Zadoks *et al.*, 1974). Spike number, the number of grains per spike, the number of grains per plant, and thousand kernel weight (TKW) were scored at harvest.

Leaf relative water content (RWC) was determined from measurements of fresh, turgid and dry weight of leaf sections from the middle part of the youngest fully expanded leaf. Turgid weight was measured after soaking the leaf sections in deionised water at 4°C over night in the dark. Dry weight of leaf sections was measured after drying at 70°C. The RWC was then calculated as 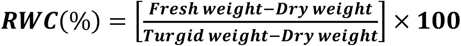 (Smart and Bingham 1974).

### 2.2 RNA extraction and gene expression analysis

Sections from the middle of the youngest fully emerged leaf were sampled for the developmental time courses at Zeitgeber Time 8 (ZT8). Samples for the diurnal expression analyses were harvested every four hours starting at ZT0, with one additional sampling at ZT22. RNA extraction, reverse transcription and qRT-PCR were performed as previously described (Campoli *et al.*, 2012*b,a*; Digel *et al.*, 2015). Several combinations of reference genes were tested for each experiment and the genes with the most stable expression were chosen for normalisation. The geometric mean of *Actin* and *ADP-ribosylation factor 1-like protein* (*ADP*) absolute expression was used for the calculation of relative gene expression levels for the developmental time courses. The geometric mean of *ADP* and *Glyceraldehyde-3-phosphate dehydrogenase* (*GAPDH*) absolute expression was used for the calculation of relative gene expression levels for the diurnal time course.

### 2.3 Statistical analysis

All statistical analyses were performed with R (R Core Team, 2020). Polynomial regressions (Loess smooth line) were calculated using second degree polynomials and an alpha of 0.75, with a 95% confidence interval. Student’s t test assuming two tailed distribution and equal variance was used to compare group means for control and drought treatments at each time point of the time course analyses with a significance cut-off of *p* < 0.05. Significant differences in trait expression between treatments and genotypes were compared by Kruskal-Wallis ANOVA followed by Conover-Iman test for multiple comparisons and Bonferroni correction with a significance cut-off of *p* < 0.05.

## 3 Results

### 3.1 Drought interacts with *Ppd-H1* to modulate flowering time

We aimed to characterise the effects of drought on the timing of reproductive development and on shoot and spike morphology. In addition, we tested if the major photoperiod response gene *Ppd-H1* controlled reproductive development in response to drought. We quantified the effects of drought on developmental timing, growth and inflorescence morphology in the spring barley genotypes Scarlett, Golden Promise and Bowman with a natural mutation in the CCT domain of *Ppd-H1* and in the derived introgression lines S42-IL107 (Scarlett), GP-fast (Golden Promise), and BW281 (Bowman) that carry wild-type *Ppd-H1* alleles introgressed from wild barley *(H. v.* ssp. *spontaneum*) or winter barley (Supplemental Figure 1) (Druka *et al.*, 2011; Schmalenbach *et al.*, 2011).

We developed an assay to apply drought starting from early vegetative growth and lasting until maturity. With this assay drought effects on the transition of vegetative to reproductive development and on floral progression were examined. Heading date, scored as a proxy for flowering time, was significantly delayed in all parental spring barley genotypes. Heading date was delayed by eleven days in Scarlett, by 13 days in Golden Promise and three days in Bowman under drought compared to control conditions (Figure 1 A). In contrast, heading date was not significantly different under drought compared to control conditions in S42-IL107 and GP-fast, and significantly accelerated in BW281. In the parental genotypes, the number of spikes per plant was strongly reduced under drought, all plants produced only a maximum of three spikes under drought compared to more than ten spikes under control conditions (Figure 1 B). The introgression lines produced on average 5–6 spikes per plants under drought compared to twice as many under control conditions and thus significantly more under drought compared to the parental genotypes. Drought also reduced the number of grains per spike in all genotypes (Figure 1 C). However, there were no consistent differences in the reduction of grain number between *Ppd-H1* variants. The reductions in the number of spikes per plant and grains per spike resulted in a severely reduced number of grains per plant under drought (Figure 1 D). Total grain numbers under drought were significantly higher in the introgression lines S42-IL107 and BW281 than the parental lines and not significantly different between Golden Promise and GP-fast. Drought did not strongly influence the TKW. Total yield per genotype was therefore primarily determined by the grain number (Figure 1 E).

**Figure 1.**
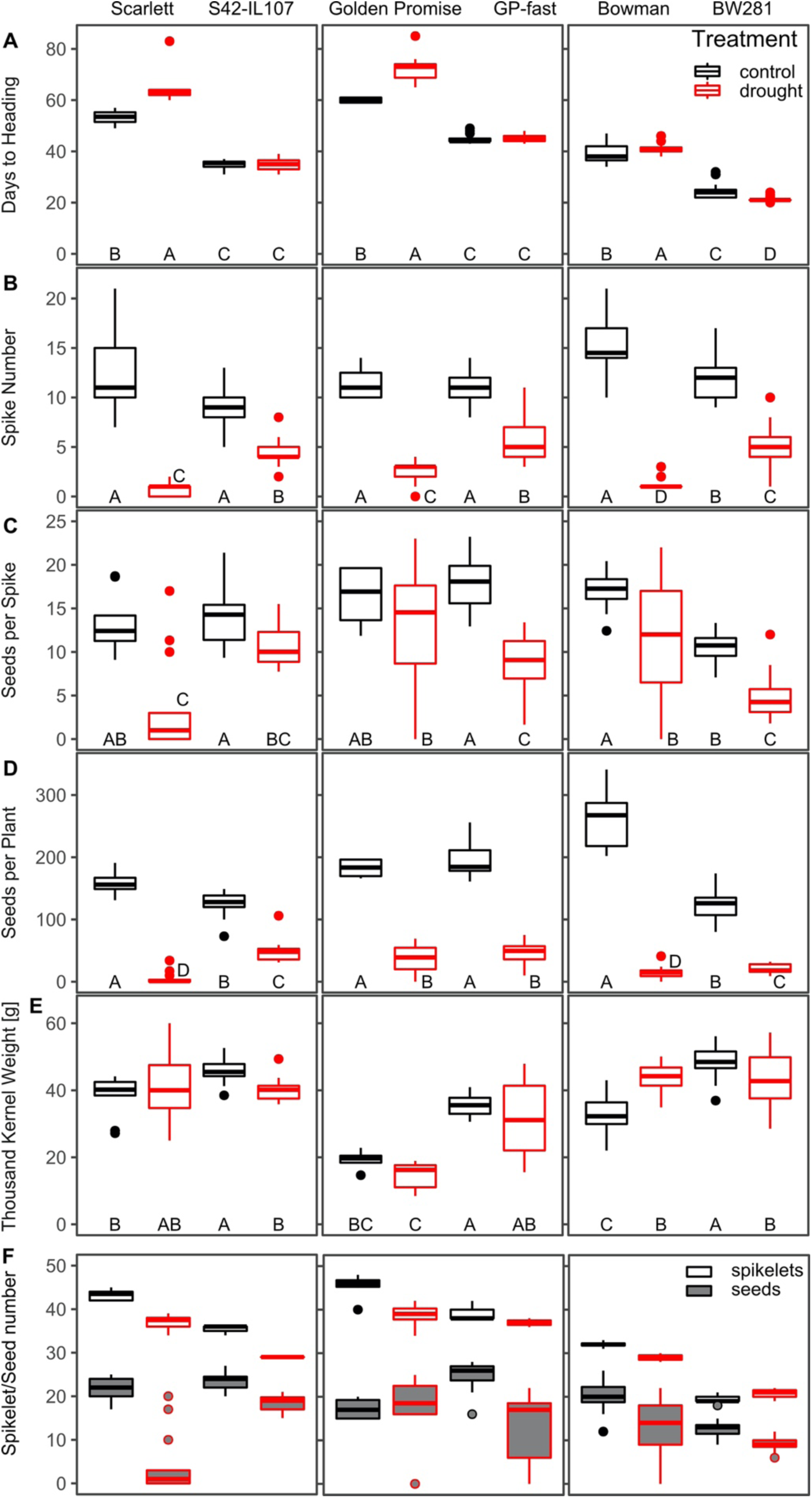
Continuous drought affects heading date, shoot and spike morphology in barley. Days to heading (A), spike number per plant (B), grain number per spike (C), the number of grains per plant (D), thousand kernel weight (TKW) (E) and the maximum number of developed spikelets (unfilled boxes) and the number of grains (grey box) (F) were scored under control (black) and drought (red) conditions under LDs (16 h light/8 h night) in the spring barley cultivars Scarlett, Bowman and Golden Promise, and the derived introgression lines S42-IL107, BW281 and GP-fast. Statistical groups were assigned using Kruskal-Wallis ANOVA and post-hoc Conover-Iman-test and Bonferroni correction. Different letters indicate groups differ (*p* < 0.05).

We further investigated at which stage drought reduced final grain number and evaluated the effects of drought on spikelet versus grain number. Drought reduced the number of initiated spikelets in Scarlett, S42-IL107, Golden Promise and Bowman by between 9% in Bowman to 18% in S42-IL107, while spikelet numbers were not significantly different between control and drought in BW281 and GP-fast (Figure 1 F). Furthermore, not all spikelets on the main spike developed grains. Under control conditions the number of grains compared to initiated spikelets was reduced by 34-37% in the introgression lines and by 37% in Bowman, 50% in Scarlett and 62% in Golden Promise. Consequently, in S42-IL107, GP-fast and BW281 a higher percentage of spikelets developed grains compared to Scarlett, Golden Promise and Bowman, respectively. Under drought conditions, the number of grains per spikelet was even more strongly reduced in all genotypes compared to control conditions, except for Golden Promise and S42-IL107. Under drought, relative grain number compared to spikelet numbers were reduced by 88% in Scarlett, by 64% in GP-fast, and by 56% and 57% in Bowman and BW281, respectively. Consequently, the reduction in grain number per spike under drought was primarily caused by an abortion of florets or floret sterility rather than a decrease in spikelet numbers.

Development of the MSA was scored after microdissection according to the scale established by Waddington *et al.*, (1983). The timing of spikelet initiation was not significantly altered by drought in any of the genotypes (Figure 2 A). However, drought delayed floral progression in the parental genotypes, but not in the introgression lines. Similarly, stem elongation, measured as plant height, was strongly reduced under drought in the three parental genotypes, but was less affected in the introgression lines (Figure 2 B). Variation at *Ppd-H1* and drought also had strong effects on the progression of tiller development (Figure 2 C). The introgression lines developed significantly fewer tillers than the parental lines under control and drought conditions. Drought delayed the development of tillers in Scarlett, Bowman and BW281, but tiller development was not significantly different in S42-IL107, Golden Promise and GP-fast. Consequently, drought had a much stronger effect on spike number than tiller number, demonstrating that the plants produced tillers during drought that did not develop a spike (Figure 1 B). The faster reproductive development in the introgression lines correlated with a reduced biomass accumulation compared to the parental lines under control and drought conditions. Drought reduced fresh weight biomass in all lines, and the relative reductions were similar between the parental genotypes and their respective introgression line. For example, 34 days after emergence a ≈70% reduction in biomass was observed in both Scarlett and S42-IL107 (Figure 2 D). We did not observe any effect of drought on the phyllochron and the number of leaves on the main culm, but leaf size was strongly reduced under drought (Supplemental Figure 2 A). Leaf RWC was not altered under drought in any of the tested lines, indicating that all plants responded to the reduced water availability through a growth reduction and thus avoided tissue dehydration (Figure 2 E).

**Figure 2.**
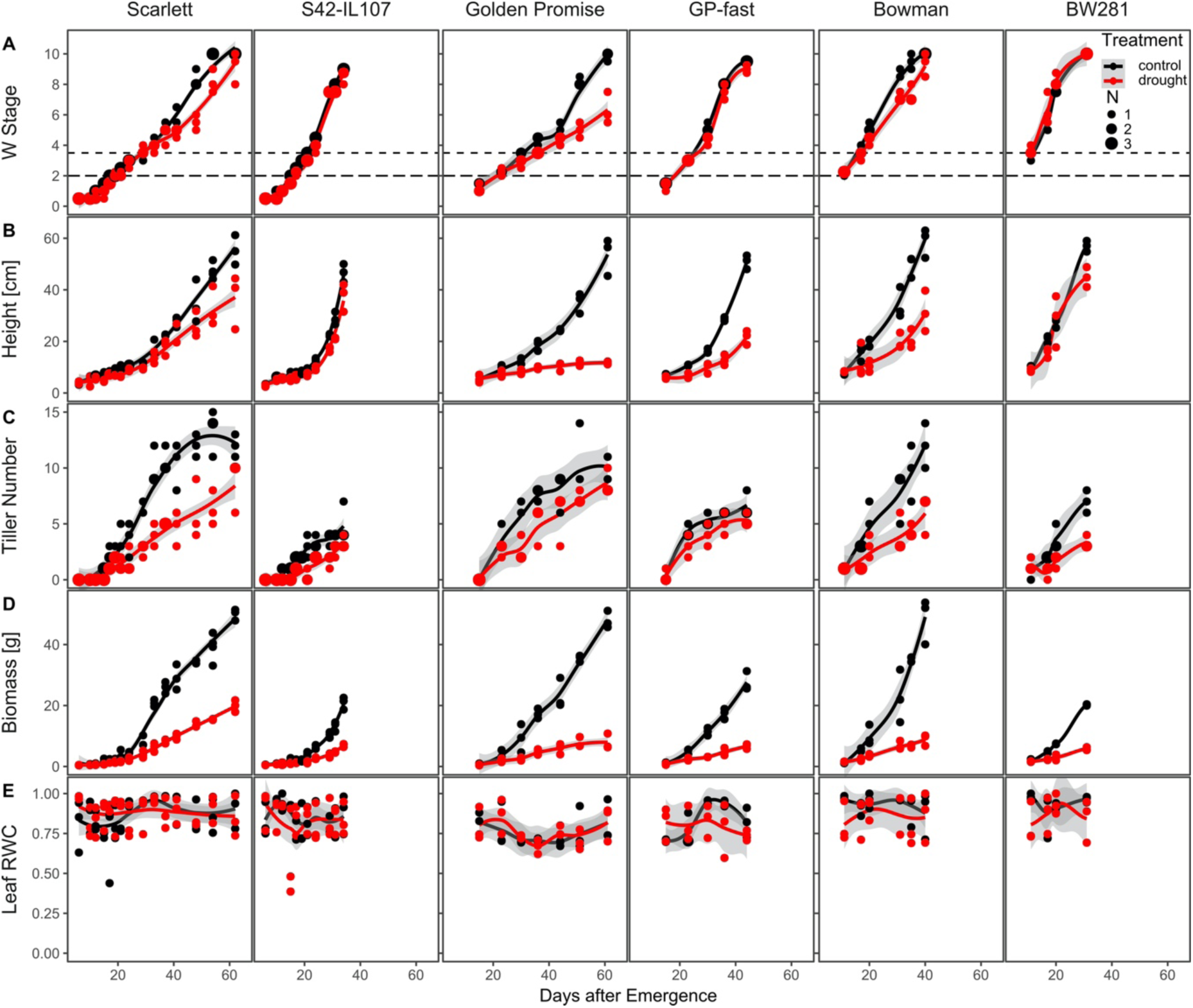
Continuous drought delays floral development in spring barley. Development of the main shoot apex (MSA) (A), plant height (B), the number of tillers (C), fresh weight biomass (D) and leaf relative water content (RWC) (E) were scored over development under control (black) and drought (red) conditions under LDs (16 h light/8 h night) in the spring barley cultivars Scarlett, Bowman and Golden Promise, and their derived introgression lines S42-IL107, BW281 and GP-fast according to the scale by Waddington *et al.*, (1983). Dot sizes indicate the number of overlapping samples. Trend lines were calculated using a polynomial regression (Loess smooth line), grey areas show 95% confidence interval.

The induction of spikelets on the MSA terminated earlier in the introgression lines which therefore formed fewer spikelets compared to their respective parents. The introgression lines initiated spikelets until W4–5 while the parental lines formed new spikelets until W5-6 (Figure 3). Under drought, the initiation of spikelets was slowed down in the parental lines, so that fewer spikelets were initiated under drought than control conditions. However, in the introgression lines, there was no significant difference in the initiation of spikelet primordia between control and drought conditions. While the parental lines initiated more spikelets than the introgression lines, a higher proportion of spikelets did not develop florets in the parental genotypes, compared to the introgression lines. The introgression lines initiated fewer spikelets under control conditions, but drought did not reduce spikelet number further in these lines. The differences between spikelet number and grain number observed in the introgression lines (Figure 1F) were therefore due to low floret fertility and not a failure in developing florets.

**Figure 3.**
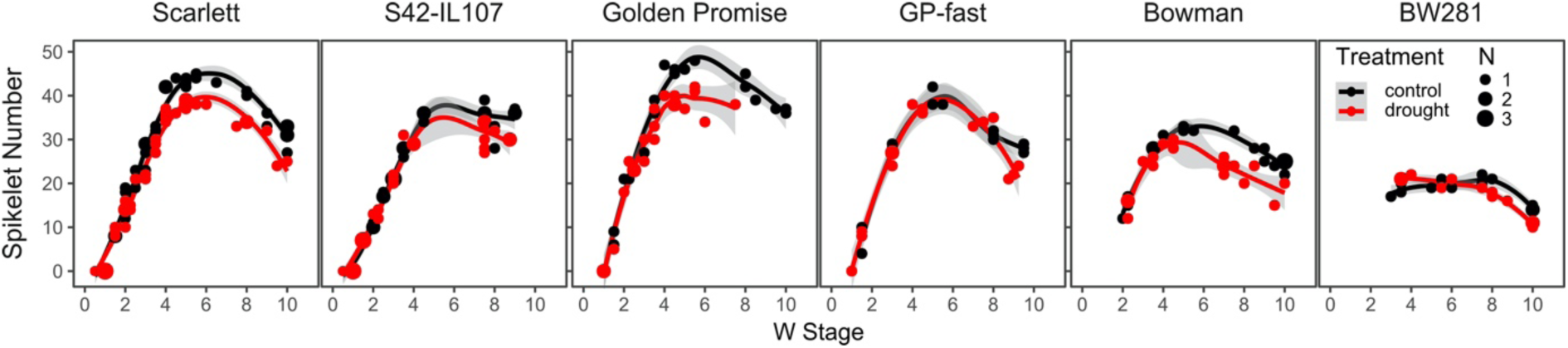
Continuous drought affects tillering and spikelet number in barley. The number of spikelets on the main shoot apex (MSA) (A) were scored over development under control (black) and drought (red) conditions under LDs (16 h light/8 h night) in the spring barley cultivars Scarlett Bowman and Golden Promise, and the derived introgression lines S42-IL107, BW281 and GP-fast. Dot sizes indicate the number of overlapping samples. Trend lines were calculated using a polynomial regression (Loess smooth line), grey areas show 95% confidence interval.

Taken together, *Ppd-H1* controlled the drought induced changes in reproductive development, shoot and spike morphology and plant height. Elite spring barley with a mutation in *ppd-H1* displayed a strong delay in floral development, reductions in plant height and the number of spikelets initiated on the main inflorescence under drought whereas these traits were scarcely affected under drought in the introgression lines with a wild-type *Ppd-H1* allele. Finally, drought had a strong detrimental effect on floret fertility which resulted in a reduction of grains independent of the *Ppd-H1* genotype.

### 3.2 *Ppd-H1* affects the plasticity of reproductive development in response to a transient drought stress

The severity, duration and timing of drought events is highly variable in nature. We therefore tested if the observed effects of drought on reproductive development are dependent on the timing and severity of the stress. In addition, we investigated if *Ppd-H1* also affected the plasticity of development in response to a transient drought stress followed by a recovery phase. Under severe drought, reproductive development stopped completely in Scarlett for the duration of the stress treatment and resumed after rewatering (Figure 4 A). However, the delay in development was maintained after the stress treatment and stressed plants flowered significantly later than control plants. In S42-IL107, reproductive development only slowed down after the onset of drought stress and did not stop completely. After rewatering, reproductive development even accelerated so that control and stressed plants flowered almost at the same time (Figure 4 A, Supplementary Figure 3). Tiller development was also halted in both genotypes upon the onset of stress but both genotypes resumed tiller development after rewatering, and tiller numbers were not significantly different between control and stress conditions at flowering. Spikelet numbers were not strongly altered during development because at the onset of drought (W3) the majority of spikelets had already initiated. Drought, however, still caused a small reduction in spikelet initiation in both genotypes. The treatment completely stopped biomass accumulation in both genotypes already after two days of withholding water. On the eight day, when the drought level was most severe, control plants of both Scarlett and S42-IL107 had accumulated almost nine times as much biomass compared to drought stressed samples. The reductions in fresh biomass were also caused by a strong decline in the leaf RWC upon application of the severe drought stress (Figure 4 A, B). However, after rewatering, RWC levels rapidly increased again and were similar to RWC levels in control plants six days after rewatering in both genotypes. While RWC levels fully recovered after rewatering and stressed plants resumed growth, fresh weight biomass was significantly lower in stressed compared to control plants at flowering.

**Figure 4.**
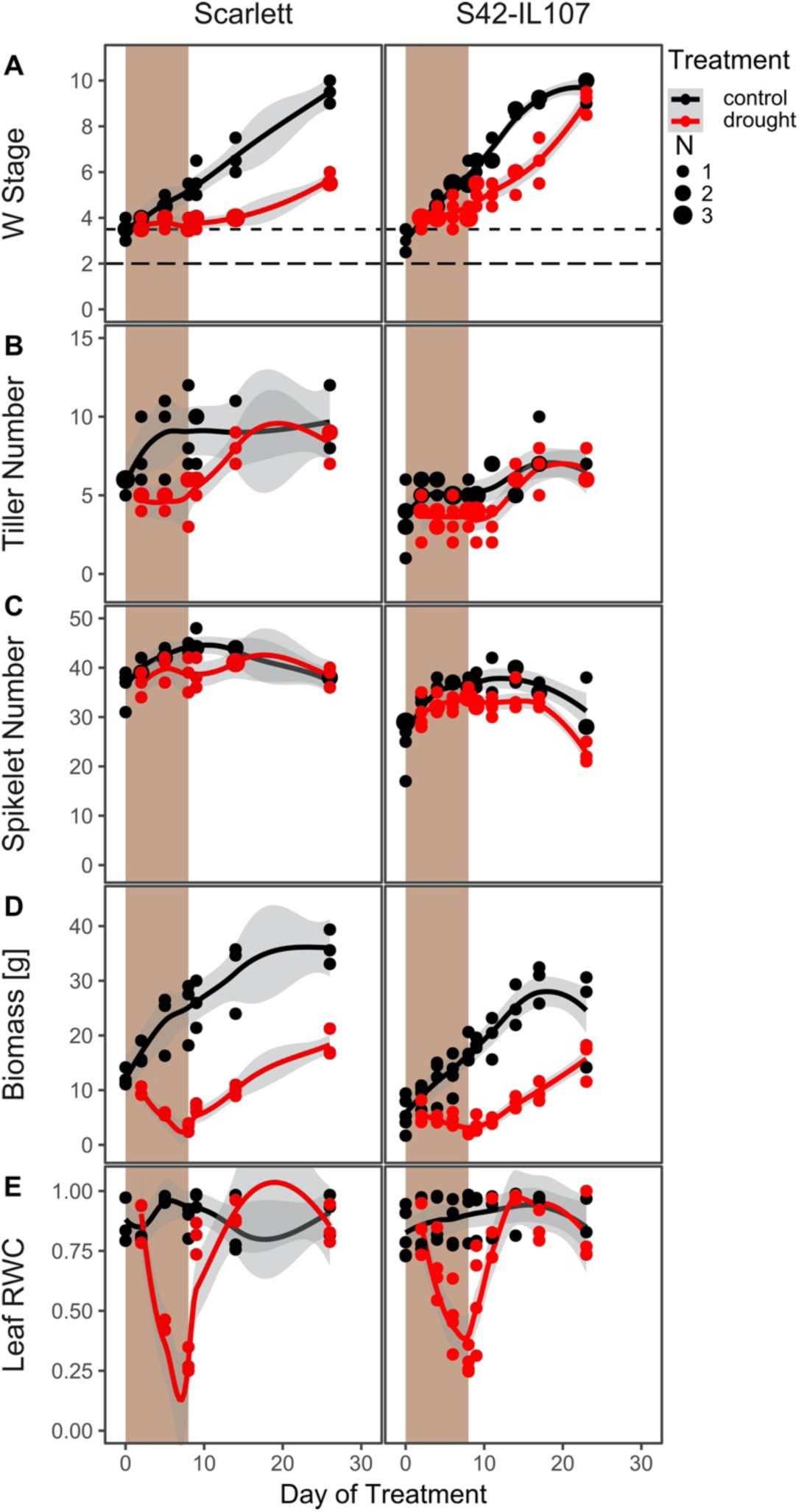
Severe drought delays MSA development in barley. MSA development (A), the number of tillers (B), the number of spikelets (C), fresh weight biomass (D) and leaf RWC (E) were scored under control (black) and severe drought (red) conditions and during recovery. Shaded areas indicate the period during which plants were not watered. Dot sizes indicate the number of overlapping samples. Trend lines were calculated using a polynomial regression (Loess smooth line), grey areas show 95% confidence interval.

Taken together, transient severe stress applied during stem elongation also delayed floral development as observed under mild stress. Interestingly, the introgression but not the parental line accelerated reproductive development after rewatering. Stressed and control S42-IL107 plants flowered nearly simultaneously, suggesting that *Ppd-H1* affects the developmental plasticity in response to drought.

### 3.3 Drought alters the expression of clock and floral regulator genes in barley

Components of the circadian clock play important roles in the control of flowering time regulators in barley. Additionally, previous studies have found that abiotic stresses alter the diurnal gene expression of core clock genes and clock regulated genes in barley (Habte *et al.*, 2014; Ford *et al.*, 2016; Ejaz and von Korff, 2017). We, therefore, examined whether reduced SWC affected reproductive development through alterations in the diurnal expression patterns of clock and flowering time genes. For this purpose, leaf samples of Scarlett and S42-IL107 plants grown under control and continuous mild drought conditions were harvested every four hours over one day of 24h at the stamen primordium stage (≥W3.5).

We investigated the expression of known barley core clock genes (Campoli *et al.*, 2012*b*; Müller *et al.*, 2019), with expression peaks at different times of the day (Figure 5). The expression levels of the morning expressed *CCA1* and the evening expressed *LUX1* were not consistently altered between drought and control conditions (Figure 5 A). Expression levels of *PRR59, PRR73* and *PRR95* and of *GIGANTEA* (*GI*) were downregulated at ZT8 under drought compared to control conditions Scarlett (Figure 5 B, C, D, F). Drought also affected the peak time of expression of some clock transcripts. The expression peaks of *PRR95* and *GI* were delayed by four hours, while expression peaks of *PRR1* and *LUX1* were advanced by four hours in both genotypes. There were no consistent differences in the expression levels and patterns of clock genes between Scarlett and S42-IL107 under both conditions. Similar to the clock genes the floral regulator genes and putative downstream targets of *Ppd-H1* were downregulated under drought. Expression of *Ppd-H1* itself was not strongly affected under drought in neither genotype (Figure 5 H). However, the expression levels of floral regulator genes differed between the genotypes under control and drought conditions. Expression levels of *FT1*, and the Barley MADS-Box genes *VRN1, BM3* and *BM8* were overall higher in S42-IL107 than in Scarlett under both conditions (Figure 5 I–L). Drought reduced *FT1* transcript levels in both genotypes, in particular at the evening peak time of expression. However, expression of *FT1* under drought was at all time points higher in S42-IL107 than Scarlett. Expression levels of *VRN1, BM3* and *BM8* were significantly higher in S42-IL107 than Scarlett under control and drought conditions. Under drought, *BM3* and *BM8* were downregulated in Scarlett at the majority of time points (Figure 5 K). In S42-IL107, transcript levels of *BM3* and *BM8* were not strongly altered between control and drought conditions.

**Figure 5.**
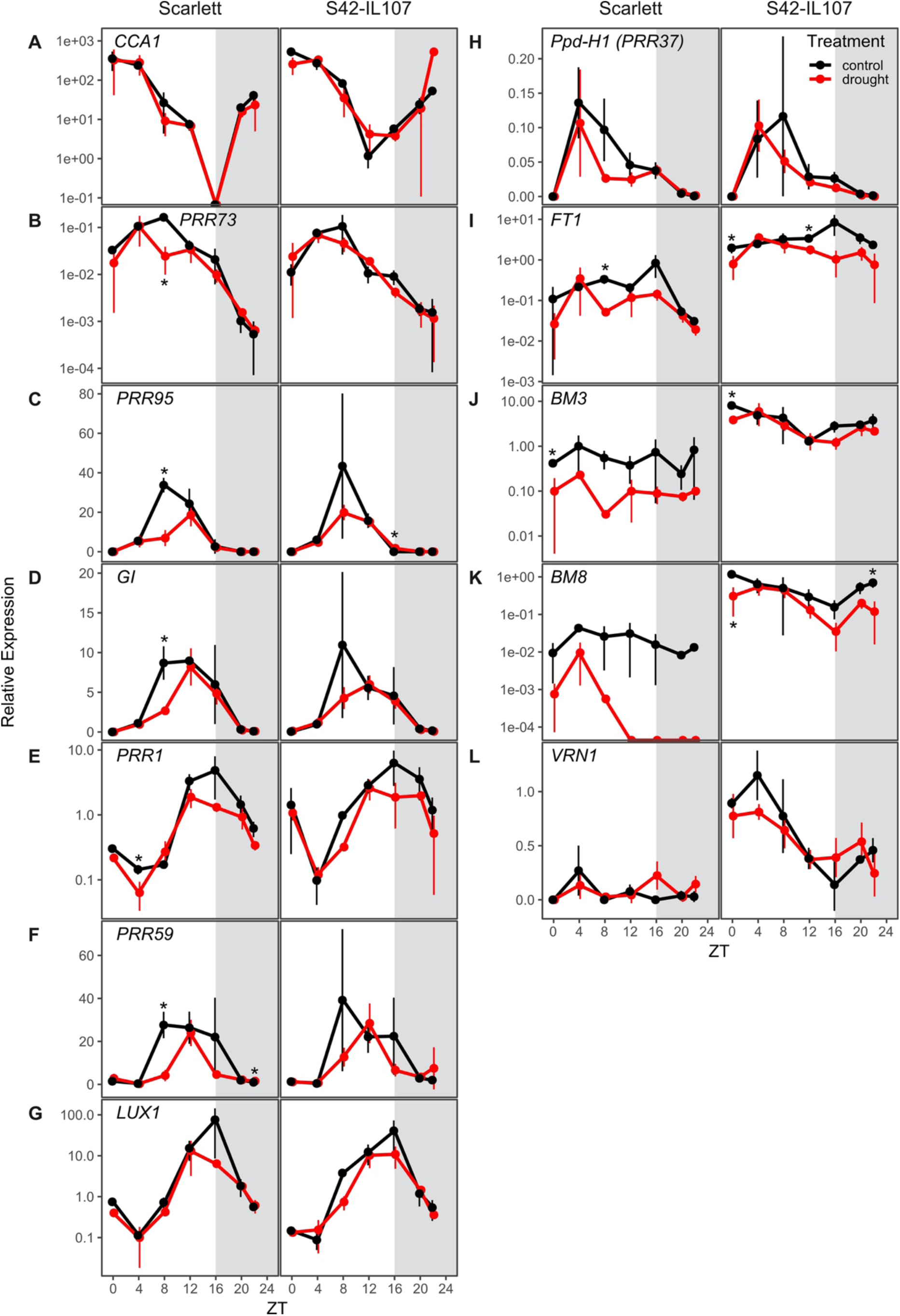
Continuous drought reduces the transcript levels of circadian clock and flowering time genes in barley. Diurnal gene expression of circadian clock genes was measured every 4 h for 24 h with an additional sampling at ZT 22 under control (black) and drought (red) conditions under LDs (16 h light/8 h night) in the spring barley cultivars Scarlett and the derived introgression line S42-IL107. Grey areas indicate darkness. Error bars indicate ± SD of three biological replicates; asterisk indicates a significant difference between control and drought at the respective time point (t-test, *p* < 0.05).

In summary, drought decreased the expression levels of clock genes and floral regulator genes and affected the peak time of expression of evening-expressed clock genes. Expression patterns of clock genes were similar between Scarlett and S42-IL107 under control and drought conditions, genetic variation at *Ppd-H1* (*PRR37*) did therefore not affect the diel expression patterns of clock genes. However, expression of floral regulator genes was significantly different between Scarlett and S42-IL107 under control and drought conditions. In addition, expression levels of floral regulator genes were more strongly altered under drought in Scarlett than S42-IL107, demonstrating that *Ppd-H1* interacted with drought to control reproductive development and expression levels of major flowering time genes in barley.

### 3.4 *Ppd-H1* alters the effect of drought on flowering time gene expression over development

We further investigated how *Ppd-H1* and drought affected expression of floral regulator genes during development in all six genotypes. Transcript levels of floral regulator genes were investigated in leaf samples from plants analysed for developmental traits as shown in Figure 2. The youngest fully developed leaf was harvested at ZT8 in all genotypes starting from the first day after transfer to LDs until flowering. At transfer to LDs all genotypes had formed a reproductive inflorescence at the double ridge stage (W2) with the exception of BW281 which was already at the awn primordium stage (W3).

The expression levels of *Ppd-H1* were not strongly altered by the treatment or *Ppd-H1* variant with exception of Golden Promise where *Ppd-H1* transcript levels were significantly higher under control than drought conditions (W3.5–W5.5) (Figure 6 A). In contrast, *FT1* expression levels were downregulated under drought in all genotypes (Figure 6 B). *FT1* transcript levels increased over development and this increase was slowed down under drought, in particular in the parental line Scarlett. In Golden Promise, no *FT1* transcript was detected under drought at any time point. In the introgression lines, *FT1* expression levels were only significantly different between conditions at single time points in S42-IL107 and GP-fast and not changed in BW281. These differences in *FT1* transcript levels under drought correlated with the observed delay in floral progression in the parental genotypes as compared to the introgression lines under drought versus control conditions (Figure 6 A). Transcript levels of the vernalisation gene *VRN1* were higher in the introgression than parental lines, but not significantly different between control and drought conditions (Figure 6 C). Transcript levels of *BM3* and *BM8* increased over development in all genotypes and this increase was delayed and reduced under drought in Scarlett, Golden Promise, and Bowman but not significantly different in S42-IL107 and GP-fast under drought versus control treatments. In BW281, *BM3* expression levels increased faster and to higher levels under drought compared to control conditions which correlated with the acceleration in floral development under drought in this line (Figure 6 D, E).

**Figure 6.**
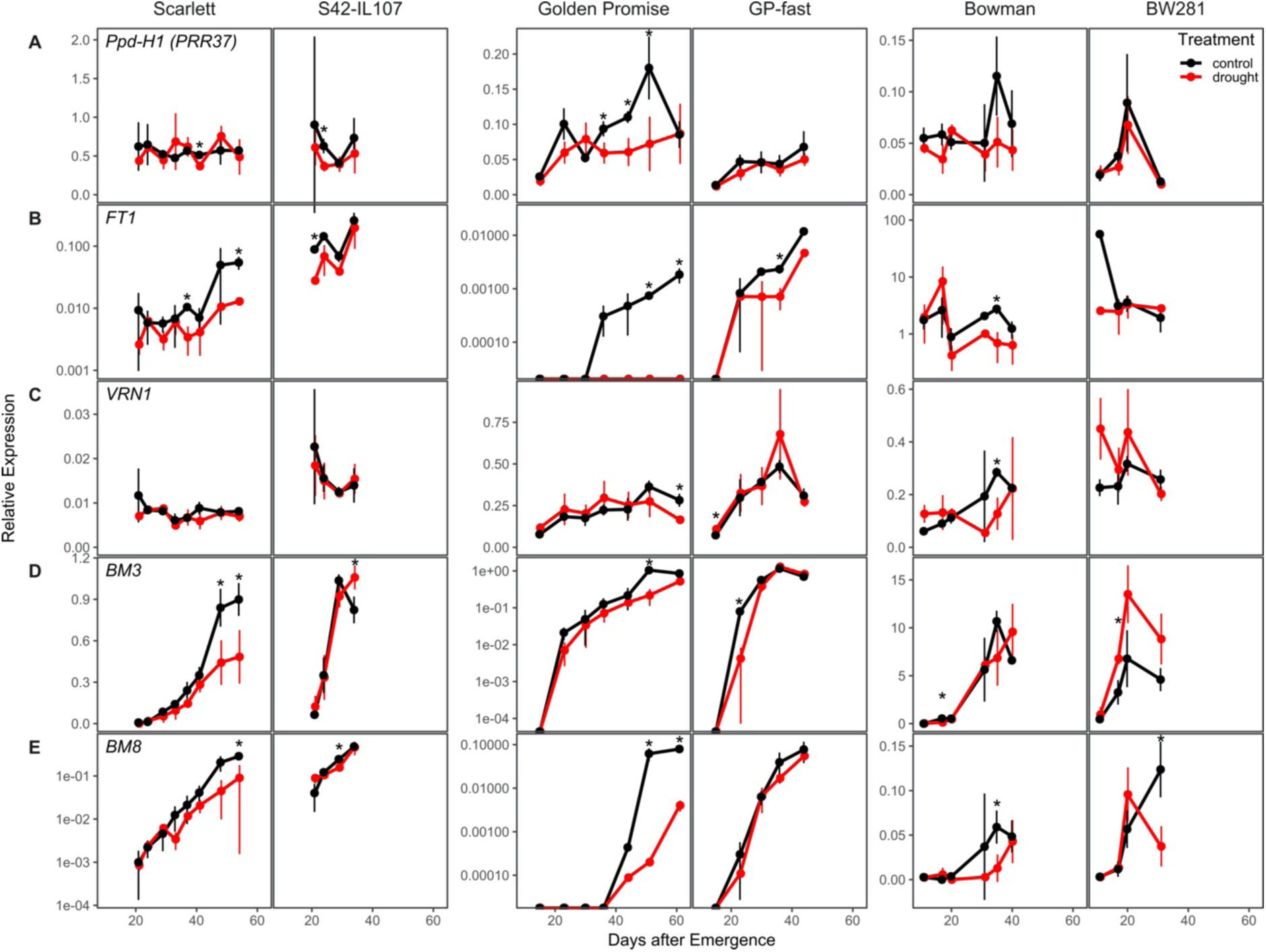
Continuous drought affects the expression of flowering time genes across development in barley. Transcript levels of flowering time genes was measured over development under control (black) and drought (red) conditions under LDs (16 h light/8 h night) in the spring barley cultivars Scarlett, Bowman and Golden Promise, and the derived introgression lines S42-IL107, BW281 and GP-fast. Error bars indicate ± SD of three biological replicates; asterisk indicates a significant difference between control and drought at the respective time point (t-test, *p* < 0.05).

We also tested the effects of the transient severe drought stress on the expression of floral regulator genes in Scarlett and S42-IL107 (Figure 7). During the transient drought treatment transcript levels of *Ppd-H1, FT1, BM3* and *BM8* were strongly downregulated compared to control conditions in both genotypes (Figure 7 A, B, D, E). In Scarlett, the downregulation of these flowering inducers extended long into the recovery phase, even after leaf RWC had returned to control levels. In S42-IL107, transcript levels of floral inducers recovered rapidly after rewatering and eventually reached the same levels as observed under control conditions. Transcript levels of *VRN1* were downregulated after the transient drought stress in both genotypes, but matched *VRN1* expression levels in control plants at flowering (Figure 7 C).

**Figure 7.**
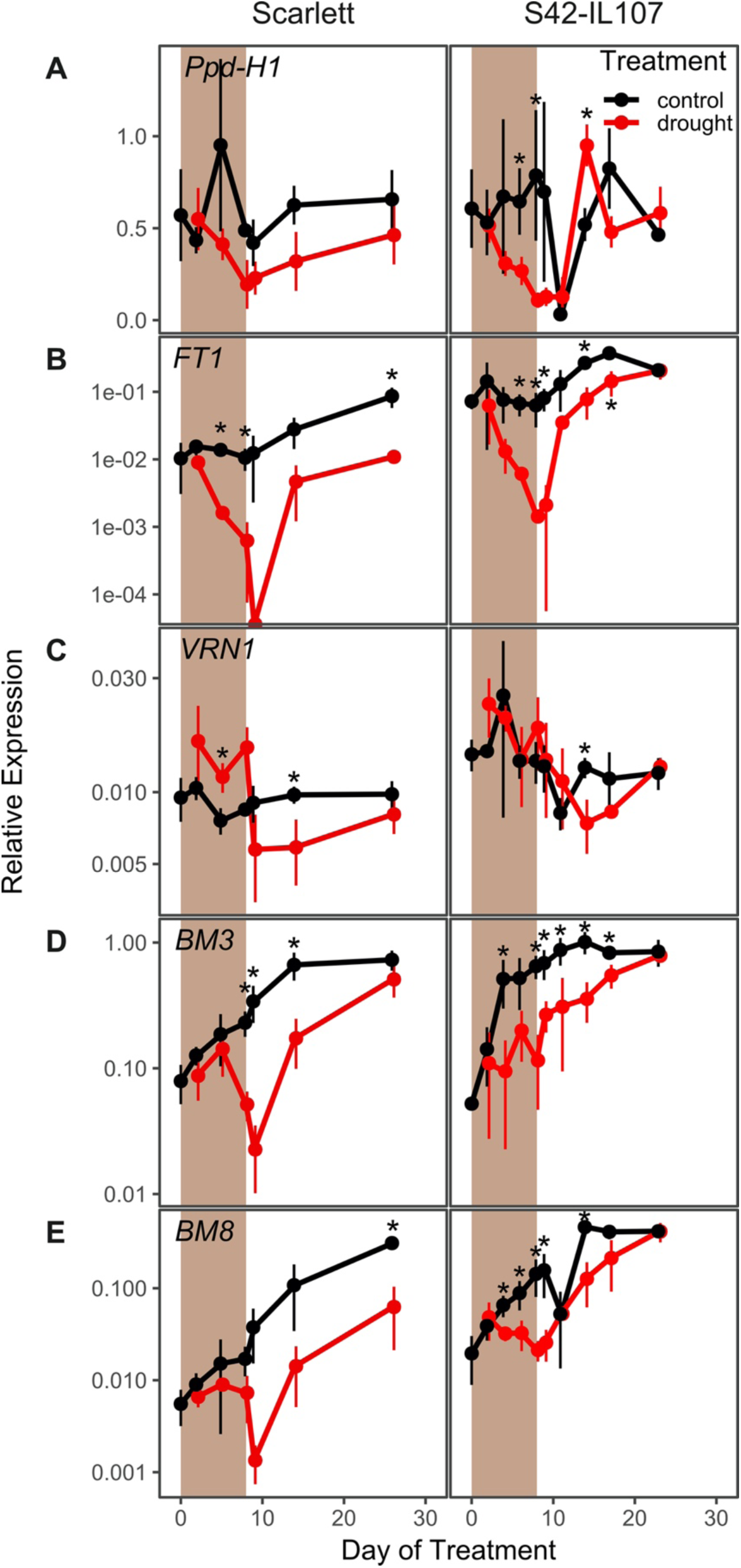
Severe drought interacts with *Ppd-H1* to control transcript levels of flowering time genes during stress and recovery. Transcript levels of flowering time genes were measured over development under control (black) and drought (red) conditions under LDs (16 h light/8 h night) in the spring barley cultivar Scarlett and the derived introgression line S42-IL107. Shaded areas indicate the period during which plants were not watered Error bars indicate ± SD of three biological replicates; asterisk indicates a significant difference between control and drought at the respective time point (t-test, *p* < 0.05).

In summary, both mild continuous and severe transient drought reduced the transcript levels of flowering inducers. However, reductions in transcript levels were stronger in the parental than the introgression lines with a wild-type *Ppd-H1* allele. *Ppd-H1* therefore modulated expression of floral inducers in response to drought in barley. In addition, transcript levels rapidly recovered after a transient drought stress to control levels in the introgression line but not the parental line, suggesting that *Ppd-H1* affected transcriptional homeostasis in response to drought.

## 4 Discussion

*Ppd-H1* was identified as a photoperiod response gene that controls adaptation to different environments by modulating flowering time in response LDs (Turner *et al.*, 2005; Cockram *et al.*, 2007; Jones *et al.*, 2008; Wiegmann *et al.*, 2019). Here, we demonstrate that *Ppd-H1* also integrates drought stress signals to modulate floral development in barley. Drought delayed floral development in the parental genotypes with a mutated *ppd-H1* allele, while reproductive development was not affected by drought in genotypes with a wild-type *Ppd-H1* allele. This variation in developmental timing in response to drought was linked to variation in the number of initiated spikelet primordia on the main shoot. Spikelet initiation was reduced in the parental lines, but not in the introgression lines under drought. Similarly, drought-triggered reductions in plant height, tiller and spike number were more pronounced in the parental lines compared to the introgression lines. Under the severe transient stress reproductive development slowed down in all genotypes, however, upon rewatering the introgression line with a wild-type *Ppd-H1* allele accelerated development so that control and stressed plants flowered simultaneously. In contrast, parental lines flowered significantly later after a transient stress than plants under control conditions. Taken together, the results demonstrated that *Ppd-H1* interacts with drought to control the development and morphology of the shoot and spike. *Ppd-H1* has already been associated with a number of shoot and spike-related traits in barley and acts as a key gene to coordinate the development of different plant organs with reproductive timing (Digel *et al.*, 2015, 2016; Alqudah *et al.*, 2016, 2018; Ejaz and von Korff, 2017; Shaaf *et al.*, 2019; Pham *et al.*, 2019). Furthermore, our results suggested that *Ppd-H1* controlled the plasticity of reproductive development in response to drought. The parental lines with a mutation in *ppd*-*H1* displayed a high trait variance between treatments and thus developmental plasticity. By contrast, the introgression lines exhibited a higher trait stability under drought, in particular for developmental timing and spikelet initiation, while biomass reductions under drought were comparable between genotypes. The identification of genes/alleles maintaining trait stability in response to environmental perturbations is interesting for breeding genotypes with high yield stability under global climatic changes and higher frequencies of extreme weather events.

The most plastic trait under drought in all genotypes was grain number. Drought caused a minor reduction in the number of spikelet primordia and in the number of spikelets/florets, but a major reduction in the final grain number. This suggested that drought reduced grain number primarily by affecting floret fertility and tiller number. It has already been described that water deficit impairs pollen development. Altered tapetal degeneration and associated changes in nutrient provision and signalling have been identified as the primary causes for cellular defects in pollen maturation under drought stress (Saini *et al.*, 1984; Lalonde *et al.*, 1997; Saini, 1997; Saini and Westgate, 1999; Aloni *et al.*, 2001; Pressman *et al.*, 2002; Barnabás *et al.*, 2007; De Storme and Geelen, 2014). Moreover, drought interferes with ovary survival or early grain development, potentially by restricting expansive growth, and thereby reduces the number of grains per spike (Oury *et al.*, 2016*a,b*; Guo *et al.*, 2016; Turc and Tardieu, 2018). Contrary to grain number, TKW was not very variable between drought and control conditions. We concluded that floral development is most susceptible to drought, while spikelet initiation as well as grain filling were less affected. This corresponds to observations in field grown wheat, where yield differences between environments were primarily controlled by variation in grain number while TKW was relatively stable across environments (Slafer *et al.*, 2014, 2015). Our results, therefore, underline the importance of floral development and fertility for yield under drought, which supports recent studies that challenge the central importance of “terminal drought” as the main cause for losses in cereal yield in drought prone Mediterranean regions (Savin *et al.*, 2015).

The circadian clock controls genes of the photoperiod response pathway and *Ppd-H1* itself is a barley homolog of an Arabidopsis clock genes (Faure *et al.*, 2012; Campoli *et al.*, 2013). Furthermore, the circadian clock controls stress adaptation and is itself regulated by stress cues (Liu *et al.*, 2013; Tamaru *et al.*, 2013; Habte *et al.*, 2014; Grundy *et al.*, 2015; Lee *et al.*, 2016; Ejaz and von Korff, 2017; Guadagno *et al.*, 2018). We consequently tested if drought interacted with variation at *Ppd-H1* to affect the expression of barley clock genes. Indeed, drought affected the amplitude and phase of clock gene expression. Clock gene transcripts were downregulated under drought, however, variation at *Ppd-H1* had no consistent effects on clock genes expression, neither under control nor drought conditions. This supports earlier studies which demonstrated that the natural mutation in *ppd-H1* did not affect the expression of other barley clock homologs neither under control condition, nor under osmotic and or high temperature stress (Campoli *et al.*, 2012*b*; Habte *et al.*, 2014; Ejaz and von Korff, 2017). However, we cannot exclude that drought might have interacted with *Ppd-H1* to affect clock proteins post-transcriptionally (Más *et al.*, 2003; Kiba *et al.*, 2007). Like the clock genes, the transcripts of the flowering time genes *FT1, BM3* and *BM8* were reduced under drought during floral development. Similarly, in rice the *FT*-homologs *Hd3a* and *RFT1* were downregulated under drought stress and this correlated with a delay in floral transition under inductive SDs (Galbiati *et al.*, 2016). By contrast, in Arabidopsis, drought induces early flowering through the ABA-dependent stimulation of GI or of ABA Responsive Element Binding Factors (ABFs) that trigger *SOC1* and *FT* transcriptional activation (Riboni *et al.*, 2016; Hwang *et al.*, 2019). On the other hand, it was also shown that ABSCISIC ACID-INSENSITIVE 4 (ABI4), a key component in the ABA signalling pathway, negatively regulated floral transition by directly promoting expression of the floral repressor *FLC*. Interestingly, the barley vernalisation gene *VRN1* was not consistently altered in expression under drought, suggesting that the vernalisation response pathway is not involved in transmitting drought signals in barley.

Because negative and positive effects of drought and ABA on flowering time were observed, it was suggested that different levels of stress may elicit different developmental responses. A moderate level of drought and ABA levels may delay floral transition, allowing for flowering to occur after the stress, while a severe drought stress and high ABA levels promote flowering and drought escape to maximise reproductive success (Shu *et al.*, 2018). However, we found that both mild and severe stress resulted in a delay in flowering time. Differential responses to drought were rather genetically controlled where *Ppd-H1* controlled the drought dependent downregulation of *FT1, BM3* and *BM8* and correlated differences in reproductive development. Furthermore, after a transient drought stress *FT1, BM3* and *BM8* transcript levels recovered fast after rewatering and eventually matched those under control conditions in the introgression but not the parental line. Consequently, *Ppd-H1* also affected transcript homeostasis after a severe transient perturbation by stress. In contrast to reports from rice and Arabidopsis drought did not strongly impact the timing of spikelet initiation but slowed down and impaired floral development and fertility. *FT1, BM3* and *BM8* have already been linked to inflorescence and floral development in barley, wheat and rice (Digel *et al.*, 2015; Wu *et al.*, 2017; Callens *et al.*, 2018; Shaw *et al.*, 2019). In rice, simultaneous knockdown of *OsMADS14* (*VRN1, FUL1*), *OsMADS15* (*BM3, FUL2*) and *OsMAD18* (*BM8, FUL3*) resulted in floral reversion and the formation of lateral vegetative tillers (Kobayashi *et al.*, 2012). Similarly, triple wheat *vrn1ful2ful3* mutants formed vegetative tillers instead of spikelets on lateral meristems and displayed a reduced stem elongation (Li *et al.*, 2019). Reduced transcript levels of *FT1, BM3* and *BM8* might therefore have contributed to an impaired floral development and decreased stem elongation in the drought-stressed plants in our study. It has been shown in barley and rice, that *FT*-homologs have positive effects on GA-biosynthesis or stem responsiveness to GA and thus stem elongation (Pearce *et al.*, 2013; Gómez-Ariza *et al.*, 2019). Reduced *FT1* transcript levels might therefore have contributed to a reduction in stem elongation under drought; Golden Promise with the strongest *FT1* downregulation under drought was also characterised by the strongest reduction in plant height.

In summary, our results demonstrate that *Ppd-H1* integrates photoperiod and drought stress signals to control reproductive timing and the plasticity of shoot and spike morphology in response to drought in barley. These differential responses to drought are linked to a differential downregulation of *FT1, BM3* and *BM8* transcripts in the leaf. Future studies need to elucidate linked transcriptional changes in the inflorescences and further dissect the effects of drought on floral organ development.

## Abbreviations

ABA: Abscisic acid
FC: Field capacity
LD: Long day
MSA: Main shoot apex
RWC: Relative water content
SAM: Shoot apical meristem
SD: Short day
SWC: Soil water content

## 6 Author Contributions

LG and MvK conceived the project and planned the experiments. LG performed all experiments, except experiments with Golden Promise which were performed by LG and EBH. LG and MvK wrote the manuscript.

## 7 Supplementary Data

Supplemental Table 1: Oligonucleotides used in this study

Supplemental Data1: Genotyping of introgression lines used in this study

Supplemental Figure 1 Genetic map of *Ppd-H1* introgression in spring barley backgrounds, introgression size in centimorgan (cM) and the polymorphic markers flanking the insertions.

Supplemental Figure 2 Continuous drought affects leaf size but not the phyllochron in barley.

Supplemental Figure 3 Severe drought delays flowering in barley.

## 8 Acknowledgements

We cordially thank Kerstin Luxa, Caren Dawidson, Andrea Lossow and Thea Rütjes for excellent technical assistance. This work was funded by the Max Planck Society and the Deutsche Forschungsgemeinschaft (DFG, German Research Foundation) under Germany’s Excellence Strategy – EXC-2048/1 – Project ID: 390686111 and the GRK2064 (Water use efficiency and drought stress responses: From Arabidopsis to Barley).

